# Neutral competition explains the clonal composition of neural organoids

**DOI:** 10.1101/2021.10.06.463206

**Authors:** Florian G. Pflug, Simon Haendeler, Christopher Esk, Dominik Lindenhofer, Jürgen A. Knoblich, Arndt von Haeseler

## Abstract

Cerebral organoids model the development of the human brain and are an indispensable tool for studying neurodevelopment. Whole-organoid lineage tracing has revealed the number of progeny arising from each initial stem cell to be highly diverse, with lineage sizes ranging from one to more than 20,000 cells. This exceeds what can be explained by existing stochastic models of corticogenesis, indicating the existence of an additional source of stochasticity. We propose an explanation in terms of the SAN model in which this additional source of stochasticity is the survival time of a lineage within a long-lived population of symmetrically dividing cells under neutral competition. We demonstrate that this model explains the experimentally observed variability of lineage sizes and we derive a formula that captures the quantitative relationship between survival time and lineage size. Finally, we show that our model implies the existence of a regulatory mechanism to keeps the size of the symmetrically dividing cell population constant.

## Introduction

The development and maintenance of the tissues and organs comprising complex organisms rely on sophisticated genetic programs to coordinate the differentiation of cells in both space and time. In many cases, this “program” does not consist of fully deterministic decision chains but instead contains stochastic components; examples include stem cell homeostasis in intestinal crypts (Snippert *et al*., 2010) and recently the development of the cortex (Klingler and Jabaudon, 2020; Llorca *et al*., 2019).

During cortical development, neurons are produced (directly or indirectly) by progenitor cells in the ventricular zone called radial glial cells (RGCs). In mice, Llorca *et al*. (2019) observed the neuronal output (i.e. number of neurons produced) of individual RGCs to vary by about one to two orders of magnitude between seemingly identical progenitors which lead them to suggest a stochastic model of cortical neurogenesis (Llorca *et al*., 2019). In human cerebral organoids (Lancaster *et al*., 2017), Esk *et al*. (2020) used comprehensive whole-organoid lineage tracing to measure the contribution of each ancestral stem cell to an organoid, and after 40 days of growth found total number of offspring to vary over four to five orders of magnitude between individual stem cells (Figure 1A). This greatly exceeds what can be explained by the stochastic model of corticogenesis of Llorca *et al*. (2019) alone, even if differences between the two model systems are taken into account. We are thus faced with the question: What causes lineage sizes in cerebral organoids to vary over multiple orders of magnitude, and how does the source of these variations fit into what is known about neurogenesis?

**Figure 1.**
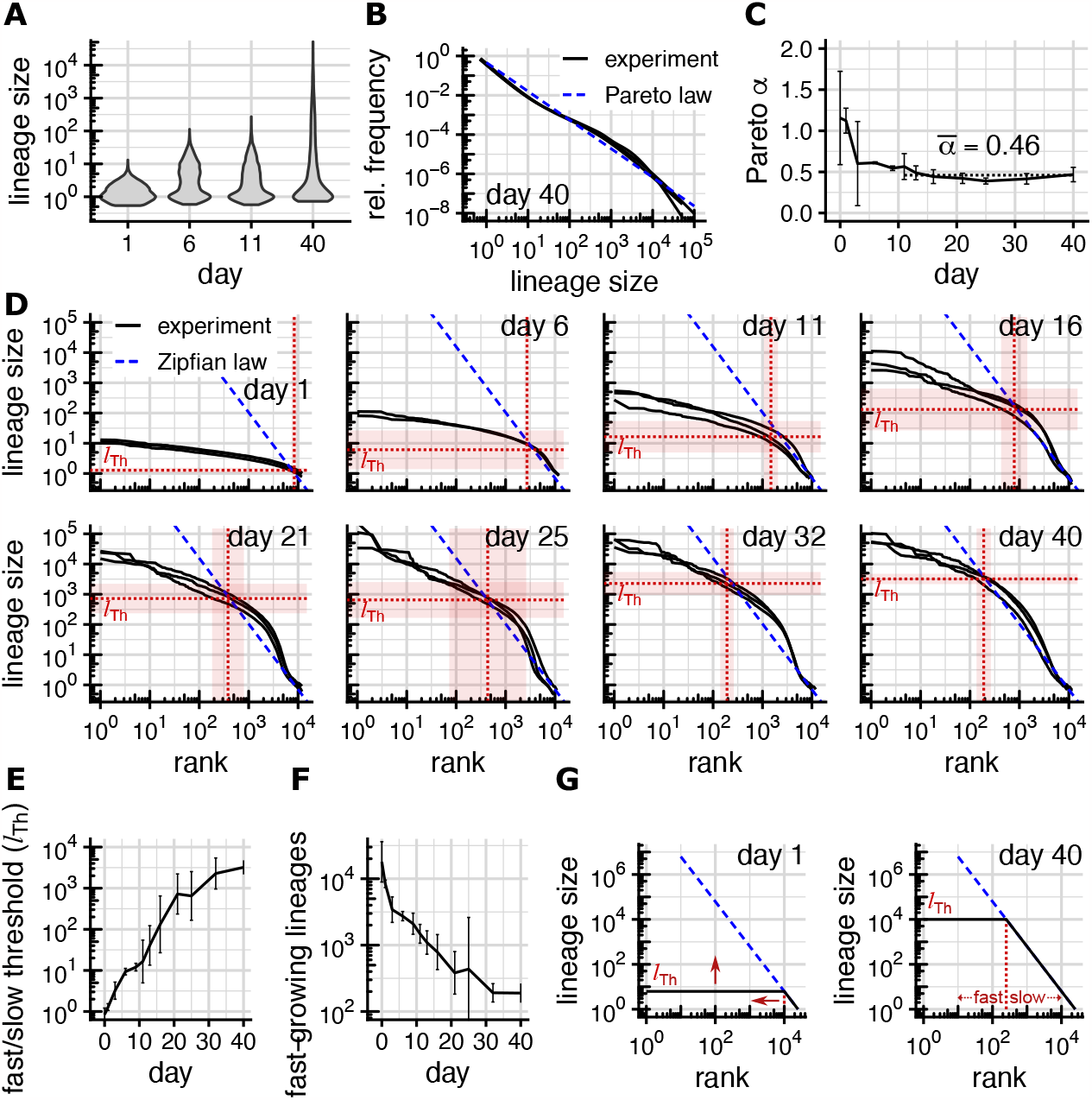
Lineages switch from fast to slow growth. (experimental data from Esk et al., 2020, error bars and shaded areas show ± two standard deviations across the three replicates) **(A)** Observed lineage size distributions at different time points (only replicate 1 shown, others are similar). **(B)** Relative frequencies of different lineage sizes on day 40 vs. Pareto power law with equality index 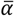. **(C)** Convergence of Pareto equality index to α = 0.46 (dotted line, represents the average from day 11 onward). **(D)** Observed rank-size distributions, threshold size s_Th_ (horizontal dotted line) and threshold rank (vertical dotted line). Threshold size and rank mark the truncation point of the Zipfian power law r^-1/α^ separating fast- and slow-growing lineages. **(E)** Size threshold s_Th_ and **(F)** number of fast-growing lineages (threshold rank l_Th_) over time. **(G)** Over time, the size of fast-growing lineages grows but their number drops, causing the lineage size distribution to approach a power law.

In the past, hidden variables like differing transcriptional state within a seemingly homogenous population of progenitors have been suggested as a possible non-stochastic source of varying offspring numbers (Zechner *et al*., 2020). For partially developed tissued with defined spatial structure and differing ancestries of cells, the existence of such hidden variables seems plausible. For cerebral organoids grown from a homogenous population of stem cells, however, differences between the internal states of cells extensive enough to explain multiple orders of variations between offspring numbers seem unlikely. Instead, we expect there to be a hitherto unobserved stochastic source of offspring variability.

Here, we show that neutral competition (also called neutral drift) within a long-lived population of roughly 10,000 symmetrically dividing stem cells we call S-cells explains the observed variations in lineage size over multiple orders of magnitude. Neutral competition has previously been shown to shape the clonal composition of tissues in homeostasis (Snippert *et al*., 2010; Corominas-Murtra *et al*., 2020), and to accurately predict the time until monoclonality (the time until all but a single lineage has died out). We show that in growing tissue like cerebral organoids, neutral competition does not lead to eventual monoclonality. Instead, the tissue’s clonal composition represents a record of the S-cell population’s history, and size differences between individual lineages reflects differences between their fates. To quantify this effect and its dependence on the size of the S-cell population, we introduce the stochastic SAN model and show that neutral competition within its S-cell population suffices to explain the observed variation of lineage sizes over four to five orders of magnitude.

## Results

### Empirical lineage size distribution

In the experiment conducted by Esk *et al*. (2020), cerebral organoids were grown from roughly 24,000 stem cells, genetically identical except for a distinct genetic barcode in each cell serving as a *lineage identifier* (LID). To determine the contribution of each initial stem cell to organoids of different ages, organoids were subjected to amplicon high-throughput sequencing. The sequencing reads (after filtering and error-correction) corresponding to each LID were counted, and the per-LID read counts normalized to an approximate number of cells comprising each lineage (see *Methods* for details).

The resulting *lineage size distribution* (Figure 1A) shows, as expected, small and equally sized lineages for organoids harvested at day 1 (lineages sizes around 1 cell). The distribution grows more uneven until day 11 (up to 30 cells/lineage) and extends over 4 to 5 orders of magnitude (up to 100,000 cells/lineage) after 40 days.

A common mathematical model for distributions extending over multiple orders of magnitude are so-called (Pareto) power laws where the frequency of objects of size *l* or larger is proportional to *l*^−α^. Parameter α is called the (Pareto) equality index because it determines how even (large α) or uneven (small α) object sizes are distributed. In double-logarithmic frequency vs. size plots, power laws appear as straight lines with slope α, which we find matches the lineage size distribution on day 40 well for α ≈ 0.46 (Figure 1B). We remark that α ≈ 0.46 represents a small equality index (i.e. diverse lineage sizes); in applications of Pareto distributions values of α often lie between 1 and 2 (Pinto *et al*. 2012).

At early time points, technical/experimental noise dominates over actual variations in linage size, and we consequently find large sample-to-sample variations of the equality index (Figure 1C) between the three replicates for each time point. Starting with day 11, all samples shown an equality index close to *α* ≈ 0.46 (Figures 1C and S2). During the same time, however, the unevenness of the linage size distribution grows considerably (Figure 1A).

### Truncated Zipfian rank-size distribution

To understand why the equality index converges while the inequality of lineage sizes grows we rank lineages by size (largest lineage first) and plot the resulting *rank-size distributions* in a double-logarithmic plot (Figure 1D). In such a plot, lineage sizes governed by a Pareto law with index *α* would be expected to be governed by a Zipfian power law (i.e. lineage sizes decrease proportional to *r*^− 1/α^ with increasing rank *r*) and because of the double-logarithmic nature of the plot would form a straight line with slope −1/*α* (Adamic and Huberman, 2002).

We observe that even from days 11 onwards (where we found *α* ≈ 0.46), this only holds for linages up to a certain lineage size threshold *l*_Th_. Above that size, lineages are multiple orders of magnitudes smaller and more uniform than the Zipfian law would predict (Figures 1D and S2). For most time points (very early ones excluded), the majority of lineages lie below the threshold and thus in the Zipfian regime. The threshold *l*_Th_ itself grows rapidly with time, increasing more than 1,000-fold (Figure 1E) over 40 days. At the same time the ratio between threshold size and largest lineage size grows only by a factor of about two (from 8.5 to 21; Figure 1D). We conclude that lineages outside the Zipfian regime grow quickly (since the threshold grows rapidly) and roughly similarly fast (since the spread of linage sizes in this regime increases only slowly). Lineages in the Zipfian, in contrast, show no overall shift towards larger lineages sizes over time, indicating that growth has mostly ceased for these lineages.

### Lineages switch growth regime

The threshold *l*_Th_ thus partitions lineages according to their growth regime into *fast*-*growing* and *slow/non-growing*. Of the (on average) 10,851 lineages that contribute to the final organoid 8,389 lineages fall into the fast-growing category on day 1; but on day 11 their number has dropped to 1.496, and on day 40 only 191 (about 2%) fast-growing lineages remain (Figure 1F). Lineages thus start in a fast-growing regime, and one by one switch to a regime of slow/no growth as time progresses. The later that switch occurs for a particular lineage, the bigger it has become before its growth ceases, leading to larger and larger lineages in the slow-growing regime and consequently to *l*_Th_ increasing as time progresses. Since lineage sizes within the fast-growing regime differ much less (about 1.5 orders of magnitude; Figure 1D) than within the slow-growing regime (3.5 orders of magnitude; Figure 1D), we neglect the differences between fast-growing lineages in a simplified representation of this model. In this cartoon version of our data-derived model, at any particular time point all fast-growing lineages have the same size. When they drop out of the fast-growing regime at a random point in time at which point their size arrests (Figure 1G).

This proposed lineage-specific switching from fast to slow growth can also quantitatively reproduce the observed truncated Zipfian with *α* = 0.46. One mathematically simple model are fast-growing lineages which grow exponentially with rate *γ* and drop out of the fast-growing regime with rate *σ* = *αγ* (S1 Supplemental Methods). But while this simple example assumes an unspecified biological mechanism behind the lineage-specific growth regime switches, we show in the following that no such mechanism is in fact necessary. Instead, we show that such growth regime switches emerge naturally from a cellular model of organoid growth.

### SAN model

We now consider a simplified, yet biologically realistic stochastic cellular model of organoid growth we call the SAN model that explains the observed linage size distributions. In our model, we distinguish between three types of cells based purely on the proliferation behavior they exhibit (Figure 2A). Cells are either *symmetrically* dividing (S-cells), *asymmetrically* dividing (A-cells) or *non-dividing* (N-cells). In our model, S-cells have the ability to self-renew indefinitely through symmetric division and can thus be considered stem cells. They form the initial cell population of an organoid, and apart from dividing symmetrically they differentiate into either A- or N-cells (or are lost, i.e. removed permanently). A-cells have committed to a differentiation trajectory and produce N-cells through asymmetric division, while N-cells do not further divide.

**Figure 2.**
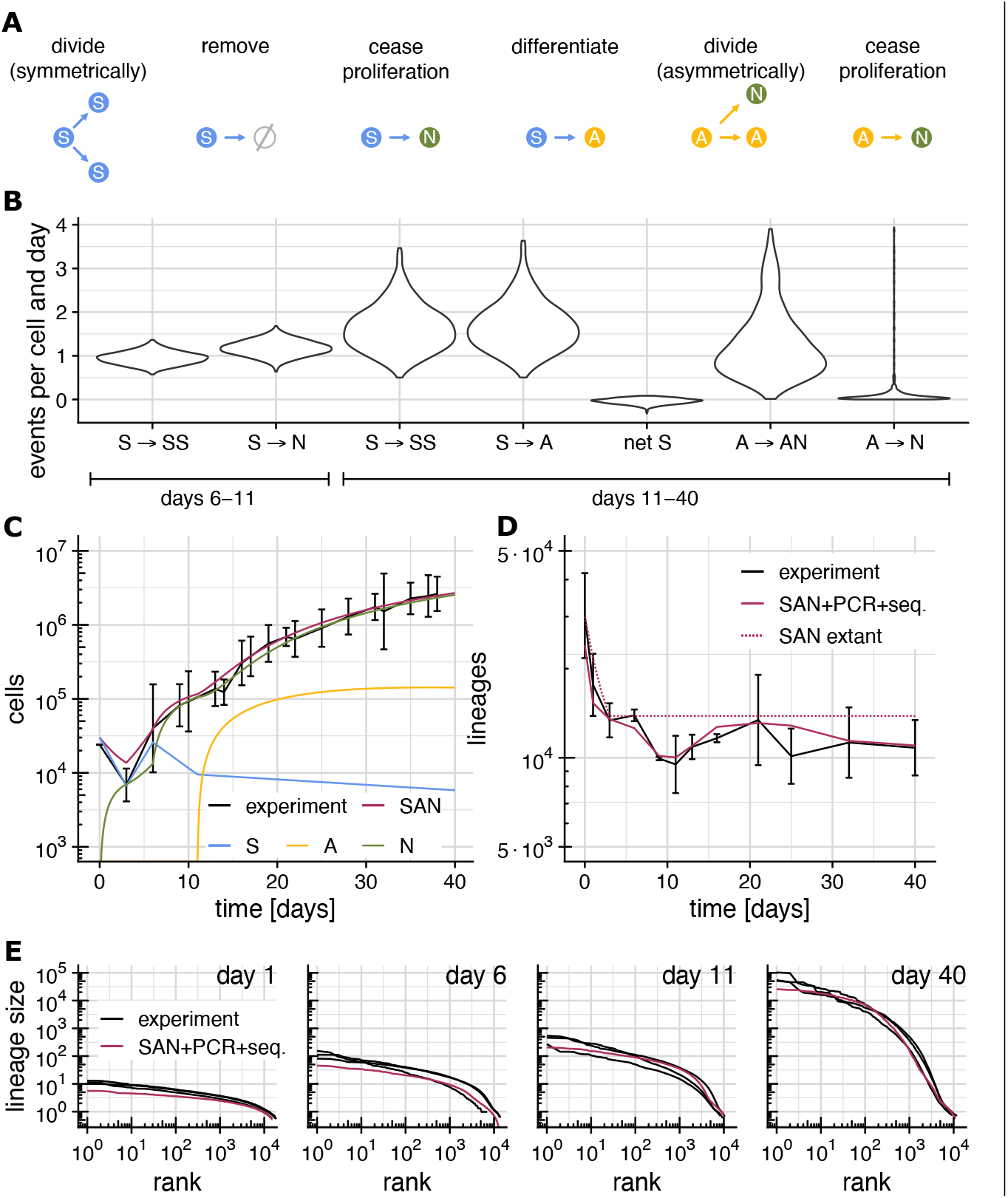
The SAN model. (experimental data from Esk et al., 2020) **(A)** Division and differentiation events that S-, A- and N-cells can undergo (see table 1 for the corresponding rates). **(B)** Posterior distribution for the estimated rates of division and differentiation events for days 6-11 and 11-40 (events not shown are assumed to have rate 0). **(C)** Total organoid size (number of cells) observed experimentally (black) and predicted by the SAN model (red), and the predicted number of S-(blue), A-(yellow) and N-cells (green). **(D)** Experimentally observed number of lineages (black), predicted number of extant lineages (dotted red) and predicted number of observed lineages (red; based on the SAN model and a model of NGS-based lineage tracing, see methods “Simulation of NGS-based lineage tracing” for details). **(E)** Rank-size distributions observed experimentally (black) and predicted by the SAN model plus a model of NGS-based lineage tracing (red).

We categorize cells into S, A and N purely based on proliferation behavior, functional categories such as RGCs do not appear in our model. We furthermore only consider such divisions as *symmetric* that produce offspring cells which are functionally identical to the parent. This precludes divisions which may appear symmetric, but where one of the offspring in fact has an altered cellular state which puts it on differentiation trajectory that leads to eventually asymmetrical division or non-proliferation. While appearing symmetric, such divisions are in the context of the SAN model more appropriately modelled as S to A transitions, since the number of eventual N-cells resulting from such a division is relatively well-determined (yet not necessarily fully deterministic.

In the SAN model, all division and differentiation events occur randomly and independently for each cell with specific time-dependent rates (table 1); from a single-lineage perspective, the SAN model is thus stochastic in nature. Any difference between the trajectories of the lineages arising from different ancestral cells is thus assumed to be purely the result of random chance, not of cell fate decisions or spatial configuration. From a whole-organoid perspective, on the other hand, the SAN model is deterministic, because random effects average out over the roughly 10,000 lineages comprising an organoid.

**Table 1.**
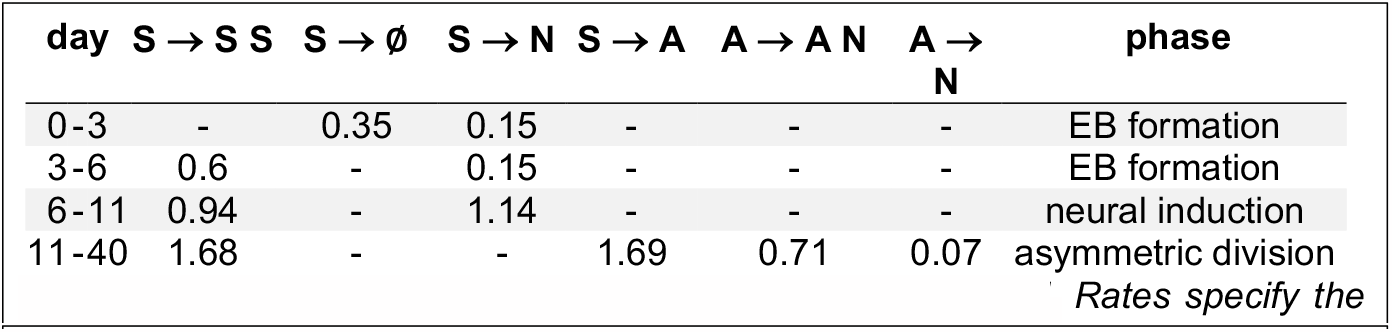
Division and conversion rates in the SAN model.

### Division and differentiation rates

To find the rates of cell division and differentiation, we split the organoid development into four time intervals (days 0-3, 3-6, 6-11 and 11-40) according to the main phases of the protocol of Lancaster *et al*. (Lancaster *et al*., 2017). Until day 6, formation of embryoid bodies (EBs) is still ongoing, and organoid development thus does not reflect development *in vivo*. For these time intervals we manually chose rates of S-cell division (S → S S), removal (S → ∅) and death (S → N). for which predicted and observed numbers of cells, lineages, and lineage sizes match experimental data (table 1). Here, distinguishing S → ∅ and S → N is necessary since dead cells within the EB are still counted by NGS-based lineage tracing, while cells which are not part of the EB are not. After day 6, EB formation is complete, and no further cells are removed from the organoid. Until day 11 S-cells are thus assumed to either divide symmetrically (S → S S) or cease proliferation (S → N), but to not produce A-cells yet. After embedding the organoids into Matrigel droplets on day 11, organoid growth enters the asymmetric division phase where S-cells are assumed to multiply (S → S S) and to differentiate into A-cells (S → A), which then produce N-cells through asymmetric division (A → A N) before they eventually cease to proliferate (A → N). We do not consider direct differentiations of S cells into N cells here since from a perspective of lineage sizes, the occurrence of such events is difficult to distinguish from slightly elevated rates of S → A and A → N. Note that A-cells decide randomly after each asymmetric division whether to continue dividing or to cease proliferation, and the N-cell output per A-cell is thus stochastic.

From day 6 onwards organoid development reflects development in vivo (Esk et al., 2020) and we hence desired to estimate the range of likely rates for each event in addition to a single most-likely value. We thus adopted a Bayesian approach comprising log-normally distributed measurement errors on top of the SAN model, and used Markov chain Monte Carlo (MCMC) sampling to find 1,000 likely rate combinations and their (posterior) probabilities (Figure 2B). To verify that the MCMC algorithm had converged to the true posterior distribution, we verified that the Gelman-Rubin diagnostic lies below 1.2 (Table S3). Finally, to arrive at a single set of most-likely values for the rates to be estimated, we then computed MAP (maximum a-posteriori) estimates (Table 1) from this posterior distribution.

Both the posterior distribution (Figure 2B) and the MAP estimates (Table 1) show the *net* rate of S-cell proliferation (the difference between the rates of S → S S and S → A) being close to zero. From this, we conclude that the size of the S-cell population changes only slowly from day 11 onwards. The posterior distributions of the other rates are, on the other hand, much broader. While the MAP estimates are thus arguably the single most likely set of rates, other combinations of rates are possible as well.

### Model validation

For the MAP rate estimates (table 1), the organoid sizes predicted between our model between day 0 and 40 agree well with the experimentally determined number of cells (Figure 2C). Similarly, the lineage size distribution predicted by the SAN model matches the observed lineage size distribution (Figure 2E). In particular, the predictions show the same truncated Zipfian distributions as the experimental data recapitulate the spread over 4 – 5 orders of magnitude. The SAN model predicts the number of *extant* lineages (lineages containing at least one S-, A or N-cell) to drop to about ≈13,700 on day 3 where it then remains. This drop in the number of extant lineages is caused by lineages that do not make it into the organoid during EB formation. This predicted number of remaining lineages (≈13,700) slightly exceeds the experimental observation (≈10,900 on day 40 on average). Since sequencing is an inherently stochastic sampling process, this excess of predicted over observed lineages caused by extant lineages which remain unobserved and is therefore to be expected. Once we take the stochastic nature of sequencing into account (see *Methods* for details) and compute the number of lineages we should expect to observe instead of the number of extant lineages, the numbers match closely (Figure 2D). To confirm the close fit of model and data, we replicated the experiments performed by Esk et al (2020) using the published protocol. These replicate experiments agree well with the original data of Esk et al. and show a similarly close fit between model and data (Figure S4).

### A-cell output

According to the MAP rate estimates for the SAN model (Table 1) a single A-cell has two options, either to continue to divide asymmetrically (probability *r*_*A*→*N*_/(*r*_*A* → *A N*_ + *r*_*A* → *N*_) = 91%) or to cease proliferation (probability *r*_*A* → *N*_ /(*r*_*A* →*A N*_ + *r*_*A* → *N*_) = 9%). Since every additional asymmetric division produces an N-cell, the likely range of additional N-cells produced over the lifetime of an A-cell is thus 0 to 30 (95% quantile; 0.91^+,^ ≈ 0.05), with an average of *r*_(*A* →*A N*_ /*r*_*A* → *N*_ = 10. This estimate number of N-cell produced by an A-cell is comparable to the range of number of neurons produced by a single RGC in mice (Llorca et al., 2019).

### Predicted S-cell population size

The MAP rate estimates (Table 1) predict that organoids contain ≈9,500 S-cells on day 11 and ≈5,800 S-cells on day 40. To take the inherent ambiguity of the MAP estimate due to the broadness of the posterior distribution into account, we computed the posterior distribution of S-cell population sizes (Figure 3A). We find that the total S-cell population size on day 11 is shows a (relatively) sharp peak at about 10,000 cells. On day 40, the estimates are more dispersed, owing to cumulative stochastic effects and to the large effect of small changes to the rates over days; yet while the exact population size is highly variable, the finding that organoids contain a significant number of S-cells on day 40 is robust.

**Figure 3.**
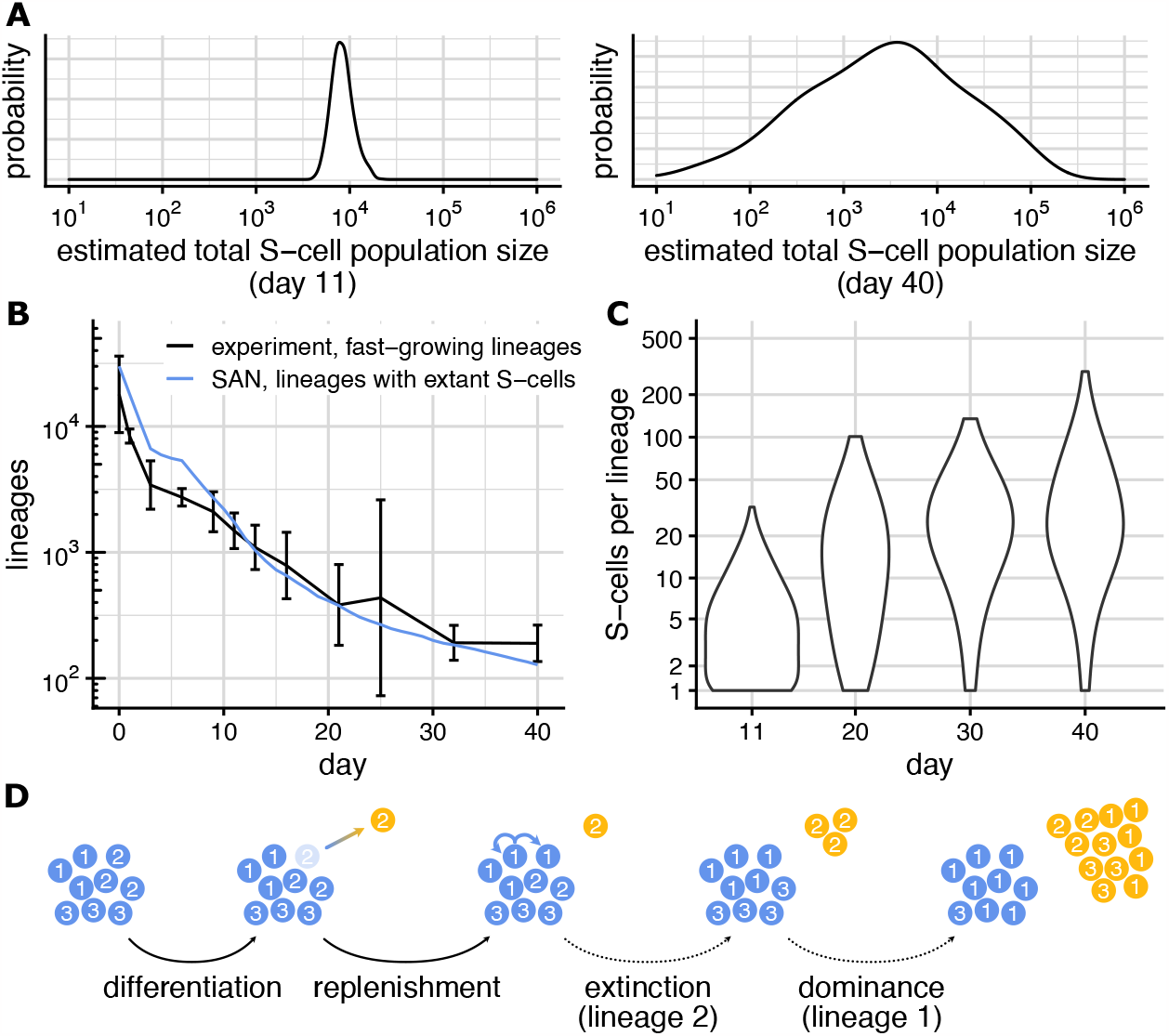
The S-cell population. **(A)** Posterior distribution of total S-cell population size on day 11 (left) and day 40 (right). Posterior distribution obtained with MCMC, see Methods. **(B)** Number of fast-growing lineages (black; from Figure 1E) number of lineages with extant S-cells as predicted by the SAN model (blue). **(C)** Distribution of S-cells per lineages (considering only lineage with extant S-cells) on days 11, 20, 30, 40 predicted by the SAN model. **(D)** Neutral competition among lineages (lineage identifiers 1,2,3) within the S-cell population. The clonal composition of the S-cell distribution changes through stochastic differentiation and replenishment events, causing some lineage to eventually lose all S-cells and others to become dominant.

### Fast-growing lineages contain S-cells

While the total size of the S-cell population changes only slowly, its clonal composition changes rapidly. From the ≈13,700 lineages comprising the organoid from day 3 forward, ≈1,700 lineages still contain S-cells on day 11, and until day 40 that number has dropped to ≈100 (Figure 3B). This drop in the number of lineages with extant S-cells is offset by an increase in the number of S-cells each of these lineages contains (Figure 3C; average grows from ≈5 cells/lineage on day 11 to ≈36 cells/lineage on day 40).

The number of lineages with extant S-cells matches the number of lineages classified as fast-growing by our fast-slow model well (Figure 3B). This implicates S-cells as being the main driver of lineage growth; as stated above (A-cell output) a single differentiating S-cells eventually on average produces 10 additional N-cells, and once a lineage contains no more S-cells its growth will thus slow down and eventually cease.

### Neutral competition shapes S-cell clonal composition

In linages with extant S-cells, the average number of S-cells grows continuously. At the same time, the spread between these lineages’ S-cells counts (from about 1 - 30 S-cells per lineage on day 11 to about 1 – 300 S-cells per lineage on day 40) grows as well. The clonal composition of an organoid’s S-cell population thus grows more and more non-uniform over time. Under the SAN model, this change is the result of *neutral competition* (Figure 3D) amongst S-cells, a term introduced to describe the population dynamics of stem cells within intestinal crypts (Snippert *et al*., 2010).

Qualitatively, the dynamics of an organoid’s S-cell population under neutral competition mimic the population-genetic Moran model (Moran, 1958) in which individuals (cells in our case) carrying different neutral alleles (lineage identifiers in our case) are randomly removed (differentiate) and are replaced (through symmetric division) by offspring of another randomly selected individual. Once the last S-cell of a particular lineage has differentiated, the lineage cannot reappear within the organoid’s S-cell population. The observed disappearance (Figure 3B) of lineages from the organoids S-cell population is thus a result of more S-cells differentiating than dividing due to random chance. Similarly, the observed growth of the remaining lineages (Figure 3C) results from more symmetric division s than differentiations, again due to random chance.

Using the SAN model, we now study the effects of neutral competition between S-cells on the clonal composition quantitatively.

### Lineage-specific S-cell extinction times determine final lineage sizes

Under the population-genetic Moran model, alleles eventually either disappear from a population or become fixed. Tissue homeostasis driven by a stem cell population under neutral competition likewise leads to eventual monoclonality, i.e. to all extant cells being eventually derived from a single ancestral stem cell (Snippert *et al*., 2010). In growing neural tissue like cerebral organoids however, the lack of constant cell turn-over restricts eventual monoclonality to S-cells. The clonal composition of the N-cell population instead records the evolution of the S-cell’s clonal composition over time; lineages whose last S-cell was lost later and/or which contained more S-cells will contribute more N-cells than lineages which die out quickly from the S-cell population.

To study the effects of S-cell extinction on lineage sizes quantitatively, we stratified simulated lineage growth trajectories according to their *S-cell extinction time* (*T*_*S*_; the time at which a particular lineage loses the last S-cell). Lineages whose S-cell population goes extinct at day *T*_*S*_ = 13 (Figure 4A left) respectively day *T*_*S*_ = 25 (Figure 4A middle) show diminished growth and a declining number of A-cells after losing their S-cells at time *T*_*S*_. In contrast, lineages whose S-cell population survives past day 40 (Figure 4A right) grow considerably faster and reach a considerably larger size. Comparing the variations in lineage sizes on day 40 between S-cell extinction time strata shows the variation due to *T*_*S*_ to dominate the variations within each stratum (Figure 4B). Thus, while other random factors have some influence, their influence on a lineage’s sizes on day 40 is negligible compared to the time the lineage loses its last S-cell.

**Figure 4.**
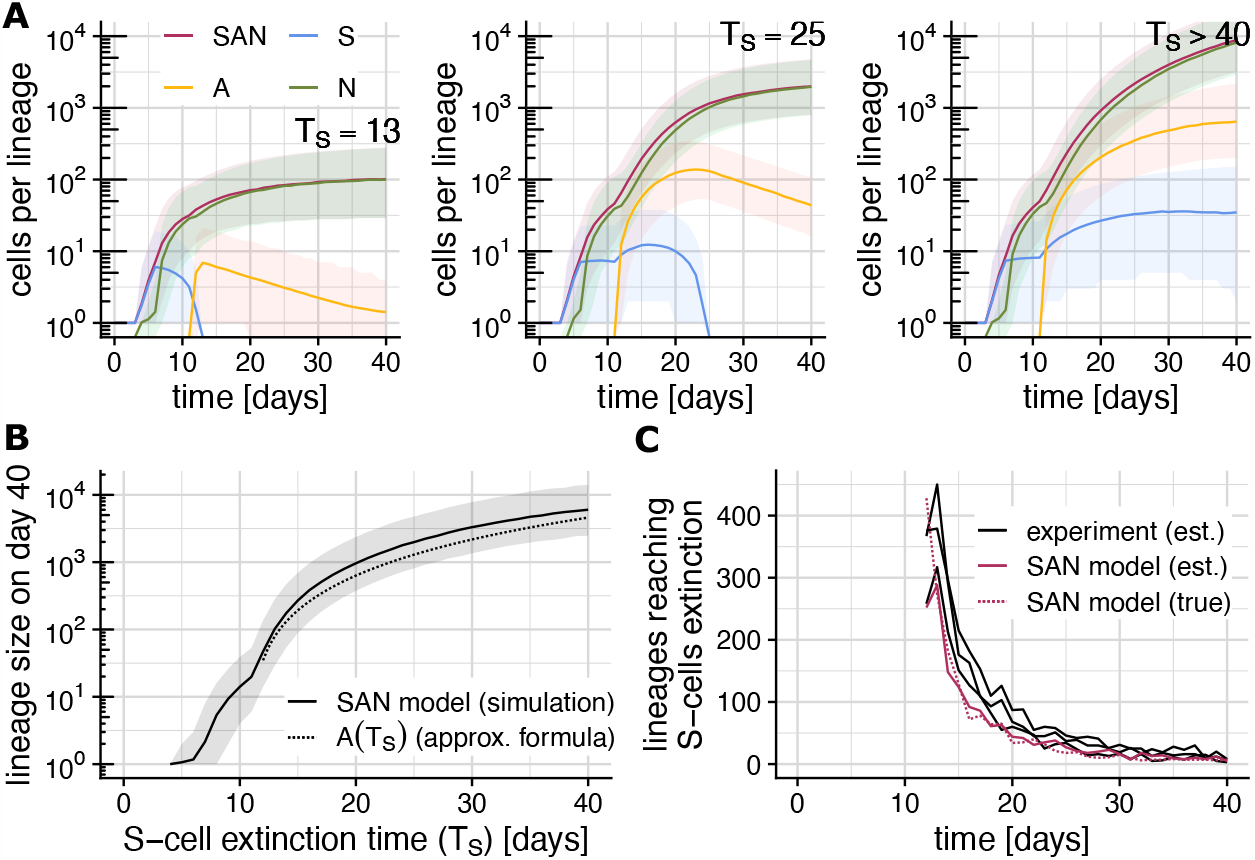
Lineage-specific S-cell extinction time determines linage size. (Shaded areas show the range between the 2.5% and 97.5% quantile across 5 billion simulations). **(A)** Lineage-specific growth trajectories under the SAN model stratified by the lineage’s S-cell extinction time T_S_. Plot show the most likely total lineage size (red), and number of S-(blue), A-(yellow) and N-(green) cells comprising the lineage. **(B)** S-cell extinction time T_S_ vs. final lineage size on day 40. Plot shows simulation results (solid) and the analytical approximation L(T_S_) (dotted). **(C)** Recovering S-cell extinction times from lineage sizes on day 40. Plot shows the number of lineages reaching S-cell extinction on each day estimated using L(T_S_) from experimental data (black) and simulated data (red), and the true number of such lineages according to the SAN model.

Mathematical analysis of the SAN model yields the approximate expression

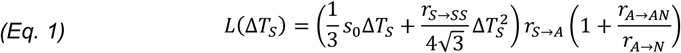

for the final lineage size of a lineage comprising *s*_0_ S-cells on day 11 (*s*_0_ ≈ 5 for rates in table 1; and we assume *r*_*S*→*SS*_ ≈ *r*_*S*→*A*_) and whose S-cell population goes extinct ∆*T s* days after day 11. We note that *final lineage size* here does not refer to the size on day 40 (or any other particular point in time), but rather to the eventual size a lineage will have reached when its growth ceases. While this does not exactly match our simulation setup (we only simulate up to day 40) the approximate final linage sizes *L*(∆*T*_*S*_) still matches the simulation results well (Figure 4B).

### Recovering S-cell extinction times from final lineage sizes

By solving the equation *L*(∆*T*_*S*_) = *L*_*i*_, the time at which a lineage lost its last S-cell can be estimated from the final size (*L*_*i*_) of that lineage. To gauge the reliability of this approach, we applied it to a simulate lineage size distribution for day 40 and found that it recovers the number of lineages that reached S-cell extinction on a particular day well (Figure 4C). When applied to the experimentally observed lineage sizes on day 40, the estimated number of lineages reaching S-cell extinction lies close to the SAN model prediction, but slightly exceeds it up to about day 30.

### Emergence of a Zipfian law

If A- and N-cells are disregarded, the SAN model is equivalent to the well-studied birth-death process. Our relationship *L*(∆*T*_*S*_) between lineage size and S-cell extinction time together with results about the distribution of S-cell extinction times ∆*T*_*S*_ (Feller, 1939) can be used to mathematically infer the lineage size distribution expected under the SAN model. We find that lineage sizes under the SAN are indeed expected to follow a (truncated) Zipfian law, but only provided that the S-cell population is long-lived and roughly constant in size. Furthermore, in this case the predicted value α = 0.5 for the Zipfian parameter closely matches the empirically observed value α ≈ 0.46 (see “Mathematical Analysis” in the methods section for details). The mathematical analysis of the SAN model thus independently supports the conclusion that S-cells divide and differentiate with very similar rates that we drew earlier from the MCMC-based rate estimates (Figure 2)

## Discussion

We have empirically observed lineages in developing cerebral organoids to initially grow fast and roughly uniformly until some stopping time, at which growth slows down significantly or ceases altogether. The stopping time is different for different lineages, and distributed such that the size of slow or non-growing lineages to follow a Zipfian power law with exponent −1/*α, α* = 0.46. While the destructive nature of NGS-based lineage tracing prevents us from directly observing lineages as they switch their growth regime, alternative hypotheses would necessarily involve either very early fate decisions, or lineage-specific proliferation rates to explain the large diversity of observed lineage sizes. Both alternative models seem unlikely given that organoids are grown from a homogenous population of stem cells.

To study the cause of the apparently random and lineage-wide switch of growth regime we consider the cellular *SAN* model. The model is intentionally coarse to keep the number of parameters tractable. Despite omitting many known details about differentiation trajectories, the SAN model still recapitulates the observed lineage size distributions well. In particular it recapitulates the experimentally observed emergence of a truncated power-law for the lineage size distribution with an α of about ½. This emergency is observed not only numerically; it also follows from our approximate expression (Eq. 1) for lineage sizes, and we recently were able to prove it using rigorous methods for a simplified version of the SAN model (Kiselev, Pflug and von Haeseler, 2023).

Under the SAN model the apparently lineage-wide switch of growth regime from fast growth to slow or no growth occurs despite the lack of either direct or indirect (e.g., through spatial colocation) lineage-wide events. Instead, growth of a lineage slows down and eventually ceases as the result of the lineage vanishing from the S-cell population through neutral competition; lineage survival time within the organoid’s S-cell population is thus what determines how long a lineage grows and is thus a major determinant of lineage size.

While neutral competition is the main source of lineage sizes variation, N-cell output of individual A-cells is also stochastic under the SAN model. This stochasticity is the result of A-cells choosing between continued asymmetric division and differentiation into an N-cell after each division. This stochasticity of N-cells produced by an A-quantitatively matches the observation of Llorca *et al*. (Llorca *et al*. 2019) that the number of neurons produced by an RGC is stochastic and varies over one to two orders of magnitude between lineages.

The relationship between a lineage’s survival times within the organoids S-cell population and the size it eventually attains can be expressed by a formula (Eq. 1). Inverting this formula allows the history of the organoids S-cell population that was recorded within its clonal composition to be read; doing so we found for days 11-30 a slight excess of lineages reaching S-cell extinction in the experimental data compared to the SAN model. We hypothesize that this might point to gradual reduction of division and differentiation rates in organoids; Since the SAN model assumes constant rates between days 11 and 40, a gradual reduction of rates would cause the model to appear to fall behind at first, and then to catch up once the true rates have fallen below the model’s rates.

While we found that we cannot estimate the rates of most division and differentiation events in the SAN model precisely, the posterior distributions of these rates indicates that the rates of S-cell division and differentiation are almost identical. The effect of these almost identical rates is that the S-cell population stays small, yet survives until day 40. Such a careful tuning of the rates of distinct events indicates the existence of an active control mechanism; without such a mechanism the rates would be expected to diverge over the course of 40 days. The simplest such mechanism may appear to be a direct, deterministic link between division and differentiation events, for example that one of the offspring of an S-cell always differentiates into an A-cell. Such a deterministic link, however, would not allow for neutral competition and thus not explain the large observed range of lineage sizes. The link between division and differentiation must thus exist on the organoid (or sub-population) level, not the level of individual cells. In the terminology of Simons and Clevers (2011), the mechanism must thus be of the *population asymmetric* type.

Given the similarity between the population-level link of S-cell division and differentiation and the dynamics within stem cell niches in intestinal crypts (Snippert *et al*., 2010), we conjecture that similar structures located within proliferation centers called *neural rosettes* (Esk *et al*., 2020) might be responsible for balancing division and differentiation of S-cells.

To study the mechanism controlling S-cell population size in more detail, it needs to be probed experimentally by perturbing organoids at specific points in time and observing their response. If a fraction of cells is killed, different mechanism would respond differently: A mechanism that relies on spatial constraints (i.e. stem cells being pushed out of a niche) would be expected to show a reduced rate of differentiations until the population has recovered. A regulatory mechanism which more directly links S-cell division to differentiation would respond differently; there we might expect the S-cell population to never reach its original size, but to instead increase its overall cell turn-over to make up for lost S-cells. We explored a few reasonable perturbations of the SAN model to see how well such perturbations can be resolved by observing the total organoid size and linage size distribution. While not all perturbations can be expected to be identifiable through their effect on the linage size distribution alone, we do observe different effects of different types of perturbations in these simulations (Figure S5).

Since the SAN model accurately predicts the lineage sizes observed for cerebral organoids grown from wildtype cells, it is also useful both when planning organoid-based perturbation screens, and when analyzing the resulting data. During the planning phase the model makes it possible to judge the effect of proliferation phenotypes on final lineage size, and thus to estimate the statistical power of different screen designs. During statistical analysis of screening data, the model provides a baseline (null model), against which the sizes of (genetically) perturbed lineages can be compared. Currently, our model and hence these applications are limited to neural organoids grown according to a protocol similar to that of Esk *et al*. (Esk *et al*., 2020). This restriction is a result of data availability. While lineage tracing data is available for a range of model systems, these datasets commonly only detected a small subset of lineages or a small subset of cells, making them unsuitable for the kind of analysis we performed in this study.

To facilitate the adoption of the SAN model, we offer an implementation of the model as an R package (http://github.com/Cibiv/SANjar).

## Acknowledgements

This work was supported by the Austrian Science Fund (FWF) project number F78.

## Methods

### Total organoid sizes

For days 0 through 21, organoid sizes were measured using fluorescence-activated cell sorting (FACS). For days 11 through 40, organoid volumes were estimated from microscopy images, and translated into cell counts using the average number of cells per volume for days 11 through 21 where both FACS and volume measurements were available.

### NGS data processing

The lineage tracing data of Esk *et al*. (2020) was obtained from GEO (accession GSE151383, supplementary file GSE151383_LT47.tsv.gz), and organoids “H9-day06-03” and “H9-day09-01” removed as outliers. Based on the assumption that in all samples the most common lineage size is 1 cell, we located the mode of the log-transformed read count distribution for every sample and used it to normalize relative lineage sizes (reads) to absolute cell counts. The validity of the underlaying assumption is confirmed by the good agreement the sum of absolute lineage sizes and the FACS and area-derived estimates of total organoid size. The normalized data is available as part of our R package *SANjar* (http://github.com/Cibiv/SANjar).

### Pareto index and fast-slow threshold estimation

For each organoid, we used the observed lineage sizes *l*_1_, …, *l*_n_ (explain n) to estimate the Pareto equality index *α* and minimal lineage size *m* with the maximum-likelihood estimator

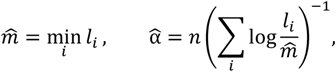

and computed the steady-state average 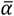 from the alpha estimates of all organoids sequenced on day 11 or later. To find the fast-slow threshold *l*_Th_ for a particular organoid, we first found intersect *d*^Pareto^ such that the Pareto-induced rank-size powerlaw 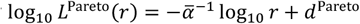 fits the size of the smallest observed lineage, and determined the smallest rank *R* for which the actual lineage size *l*_(*r*)_ matches or exceeds the power law *L*^Pareto^(*r*). We then fit a separate log-log-linear model log_10,_ *L*^Large^(*r*) = *k* log_10,_ *r* + *d*^Large^ to lineages with ranks 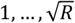 (which we assume are surely not governed by the Pareto law), and set *l*_Th_ to the size at which the two laws intersect (meaning *l*_Th_ = *L*^Pareto^(*r*) = *L*^Large^(*r*)).

## SAN model simulation

The total number of S-, A- and N-cells that an organoid is predicted to comprise at time *t* is computed based on the deterministic SAN model (with rates *r*_*S*→*SS*_, *r*_*S*→∅_, *r*_*S*→=*A*_ *r*_*S*→ *N*_, *r*_*A* →*A N*_, *r*_*A* → *N*_ of these events occurring per cell and per day). The deterministic SAN model is described by the ordinary differential equations (ODE),

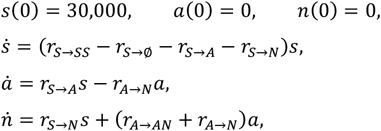

which can be solved analytically (for the time-homogenous case) and is then evaluated separately for each time interval within which rates are constant (The initial number *s*(0) of S-cells is set to 30,000 instead of 24,000 to account for a slight excess in the number of observed lineages on day 0, likely due to a combination of multiple labelling and sequencing artefacts).

To find the predicted lineage size distribution at time *t*, the stochastic SAN model is simulated independently for each of the 30,000 lineages in an organoid. The simulation proceeds in discrete time steps Δ*t*, which are chosen small enough to make the probability of a single cell undergoing two events negligible (< 10^−3^). Given the numbers *S*_*i*_ (*t*), *A*_*i*_ (*t*), *N*_*i*_ (*t*) of S-, A-, N-cells comprising lineage *i* at time *t*, the number of cells Δ_*e*_ undergoing event e is chosen from a Poisson distribution. Specifically,

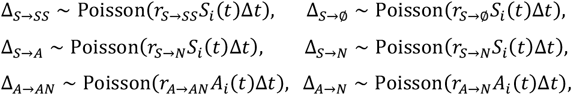

and the number of S-, A-, N-cells at time *t* + Δ*t* is then set to be

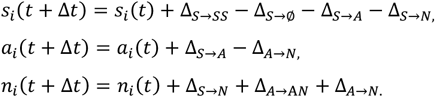

Finally, the lineage size distribution *l*_1_, …, *l*_30,000_ at time *t* is found by summing up the number of S-, A- and N-cells, *l*_*i*_ (*t*) = *s*_*i*_ (*t*) + *a*_*i*_ (*t*) + *n*_*i*_ (*t*).

While continuous-time methods exist to simulate the SAN model exactly, these methods, have the drawback that that they simulate each event separately. Their runtime thus strongly depends on the total number of cells produced, and hence on the parameter values. Since the MCMC strives to fully explore the parameter space, continuous-time algorithms perform much worse than our discrete-time algorithm, to the extent that they make running MCMC to convergence impractical. To ensure that using a discrete-time algorithm did not affect our results, we compared our some of our simulation results against an exact algorithm verified that the differences were negligible.

### Simulation of NGS-based lineage tracing

The effect of PCR amplification and sequencing on the observed lineage sizes was simulated using a stochastic model of PCR amplification and sequencing (Pflug and von Haeseler, 2018) with parameters *PCR efficiency* and average *reads per molecule* (in our case per *lineage*). For every sampling time *t*, we simulated one read count (normalized to one read per cell on average) per lineage; parameters were *PCR efficiency* 35% (estimated from the day 0 data) and average *reads per lineage Wl*_*i*_/Σ_*i*_ *l*_*i*_ for a linage comprising *l*_*i*_ cells (*W* is the median experimental library size for time *t*). The simulated read counts where then normalized to cells by division by the average number of reads per cell (*W*/Σ_*i*_ *l*_*i*_).

### SAN rate estimation

For days 0-3 and 3-6, rates which replicate the experimental data well were found by trial and error. For the remaining 6 biologically relevant rates (of S → S S and S → N between 6 and 11, and S → S S, S → A, A → A N and A → N between days 11 and 40) we computed the posterior distribution given experimentally observed total organoid sizes ĉ ^(*dayt*)^ (on days *t* ∈ 𝒟 = {0, 3, 6, 9, 10, 13, 14, 16, 17, 19, 21, 22, 25, 28, 31, 32, 35, 37, 38}) and ranked lineage sizes 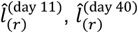(on days 11 and 40, for ranks *r* ∈ ℛ = {1, 2, 5, 10, 15, 25, 40, 60, 100, 150, 250, 400, 600, 1000, 1500, 2500, 4000, 6000, 10000, 15000, 25000}). To account for biological differences between replicates we assumed that experimental observations are log-normally distributed around the SAN model predictions *c*^(*dayt*)^ and 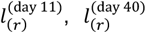; the likelihood of the rate vector *θ* (comprising the 6 rates mentioned above) given the experimental data is thus

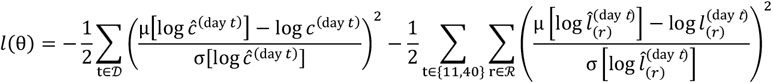

where *μ*[…] and *σ*[…] denote the mean respectively standard deviation across biological replicates. Rates were restricted to lie between 0 and 4 and *a priori* assumed to be equally probable; the posterior probability of *θ* is thus proportional to *l*(*θ*). To find this posterior distribution, we sampled 1,000 random rate vectors according to their likelihoods by simulating 1,000 Markov chains using pseudo-marginal Metropolis-Hastings Markov chain Monte Carlo sampling (Beaumont 2003; Andrieu & Roberts, 2009; Warne *et al*., 2020). Each chain was initialized with random parameters drawn uniformly from [0,4], and each chain was run until it had accepted 1000 moves. To verify convergence of the MCMC chains, we computed the Gelman-Rubin (GR) diagnostic (Gelman, 1992) for each estimated parameter (plus the net S rate, i.e. the difference between the rate of S-cell production and differentiation). Our choice of uniformly distributed initial values satisfies the requirement over being over-dispersed relative to the posterior distribution, making the GR diagnostic a applicable. For all rates except A → N, the 95% confidence intervals for the GR diagnostic lay below 1.2. For the rate A → N, the upper bound of the CI was 1.24 and the point estimate 1.19 (table S3). We concluded that the MCMC chains have explored the parameter space sufficiently deeply for the posterior distribution to be accurate. To find point estimates of the parameter, we computed the maximal mode of the (joint) posterior distribution with the mean-shift algorithm to obtain the MAP estimates (table 1).

### Mathematical Analysis

If we consider only S-cells, the SAN model corresponds to the well-known birth-death process (Feller, 1939). We consider the diffusion approximation of this process and restrict our mathematical treatment to day 11 and later where the rates of symmetric division (*r*_*S* → *S S*_; birth) and of differentiation (*r*_*S* → *A*_ ; amounts to death since we consider only S-cells) are similar enough to be considered identical (*r*_*S* → *S S*_ = *r*_*S* → *A*_ = λ/2). The number of S-cells within a lineage at time *t* (where *t* = 0 represents day 11) is then governed by the stochastic differential equation (SDE)

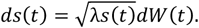

Using Onsager-Machlup theory (Onsager & Machlup, 1953; Dürr & Bach 1978) it is possible to find the most probable trajectory of a linage that contains *s*_0,_ cells at *t* = 0 and loses its last S-cell Δ*T*_*S*_ days later (Kiselev, Pflug & von Haeseler 2023),

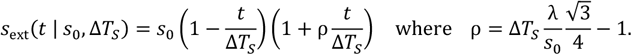

On average, a lineage grows by *r*_*S* → *A*_ A-cells per S-cell and per day, and over its lifetime every A-cell will eventually produce *r*_*A* →*A N*_ /*r*_*A* → *N*_ additional N-cells through asymmetric division. Eventually, a lineage that starts out with *s*_0_ S-cells and loses its last S-cell Δ*T s* days later will thus approximately grow to size

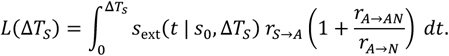

Integration of this expression and setting λ = 2*r*_*S*→ *S S*_ yields Eq. (1).

For an S-cell population governed by the SDE 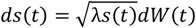, the probability that Δ*T*_*S*_≥ *t*, is exp (−2*s*_0_/λt), and for sufficiently large *t* the probability that Δ*T*_*S*_≈ *t* is thus approximately 2*s*_0_*t*^−2^ /λ. By using Eq. (1) to translate this distribution of Δ*T*_*S*_ into a distribution of lineage sizes we find that the distribution of lineage sizes with large extinction times Δ*T*_*S*_ should follow a Pareto distribution with *α* = 1/2. A more detailed mathematical analysis of the SAN model with an emphasis on criticality can be found in (Kiselev, 2022).

## Supplemental Information

**S1 Supplemental Methods. Additional details about some procedures**

**S2 Figure. Relative lineage size frequencies for days 1 through 40**.

**S3 Table. Gelman-Rubin diagnostic for MCMC results.**

**S3 Figure. Replicate Experiments**.

**S5 Figure. Perturbation Simulations**.

## S1 Supplemental Methods

### Relative linage size frequencies

To plot the empirical lineage size frequencies (Figure 1B, Figure S2) we first assigned each lineage size a density by numerical differentiation of the empirical CDF (after conversion of the ECDF from a piece-wise constant to a piece-wise linear function). The resulting density was then smoothed using loess regression with degree 2 and smooth parameter 0.75 as implemented by R’s loess function.

### Exponential growth of the size and decay of the number of fast-growing lineages

A mathematically simple example of a model that predicts lineage sizes to follow a truncated Zipfian law with index *α* is that of exponential grow (with rate *γ*) of fast-growing lineages together with an exponential decay (with rate *σ* = *αγ*) of the number of fast-growing lineages. Under this model, at any time *t* a fraction of *e*^− *σ* t^ lineages are still growing at thus have size *e*^γ t^. The fraction of lineages that have ceased growth around time 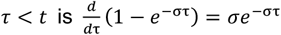, and these lineages have size *s* = *e*^γ τ^. Treating *s* = *e*^γ τ^ as a change of variables between extinction time τ and lineage size yields for lineage sizes *s* < *e* ^γ τ^ that the probability of a lineage having a size between *s* and *s* + *ds* is

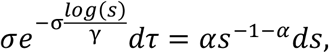

which represents a Zipfian distribution with index *α* truncated at size *e*^γ τ^.

### Perturbation Simulations

We simulated three perturbed versions of the SAN model where we modified the rates of division and/or differentiation starting with day 11. (A) A reduced rate (one half, and one tenth) of asymmetric division and therefore a reduced output of N-cells per A-cell. This can be interpreted as a model of reduced neurogenesis by precursor cells such as RGCs. (B) Reduced rate of S-cell division and differentiation while keeping the net growth rate (division minus differentiation) intact. This can be interpreted as a simple model of an ASPM knockout. (C) No symmetric divisions post day 11, rate of asymmetric division as large as is biologically realistic (2 divisions per day) and no final differentiation of A into N cells (rate of A to N differentiation is zero). Even though the N-cell output of A-cells is maximized, the lack of symmetric division reduces the maximal linages sizes by an order of magnitude.

### Replicate Experiments

Stems cells were labelled, and organoids grown and sequenced as described by Esk et al. (2020). Sequencing data will be deposited into GEO before publication.

**Figure S2.**
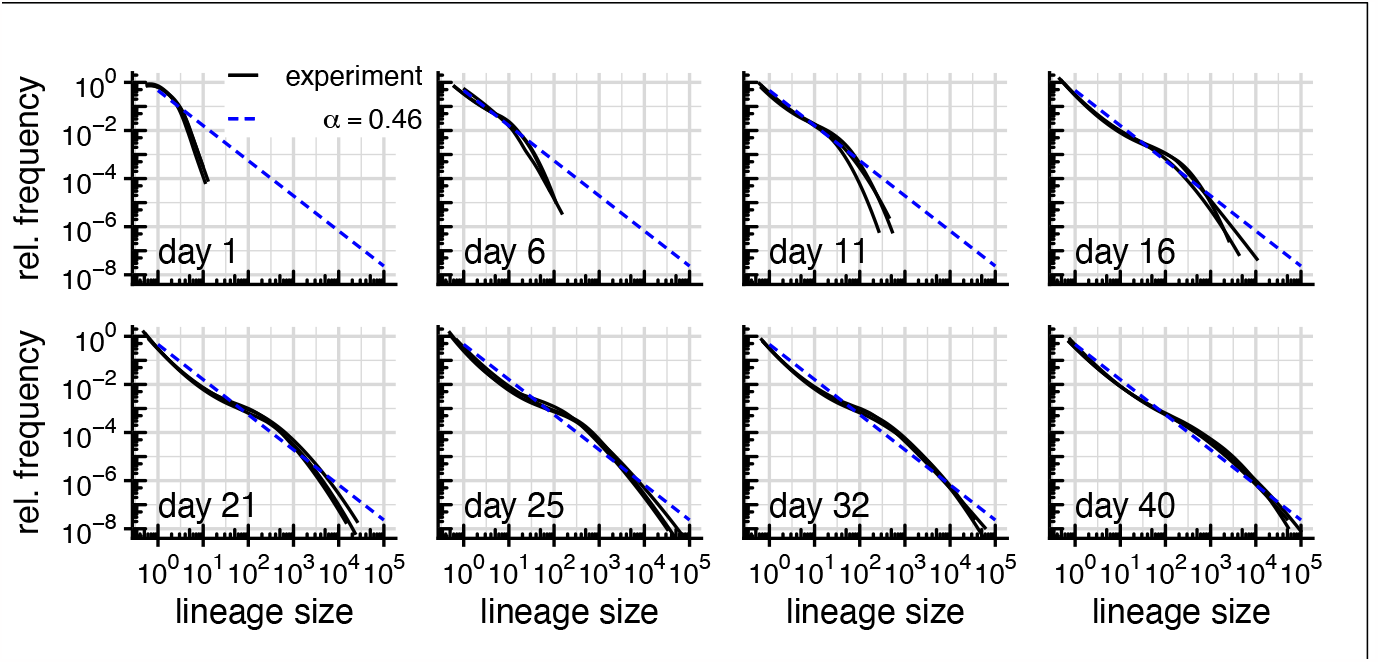
Relative lineage size frequencies for days 1 through 40. Relative frequencies of different lineage sizes on day 40 vs. Pareto power law with equality index 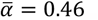.

**Table S3.** Gelman-Rubin diagnostic for MCMC results. Values of the point estimate and upper bound of a 95%-CI of the Gelman-Rubin diagnostic (Gelman, 1992).

|  | days 6 - 11 |  | days 11- 40 |  |  |  |  |
| --- | --- | --- | --- | --- | --- | --- | --- |
| GR | S → S S | S → N | S → S S | S → A | net S | A → A N | A → N |
| point est. | 1.12 | 1.14 | 1.15 | 1.15 | 1.14 | 1.15 | 1.19 |
| upper CI | 1.16 | 1.18 | 1.19 | 1.19 | 1.18 | 1.19 | 1.24 |

**Figure S4.**
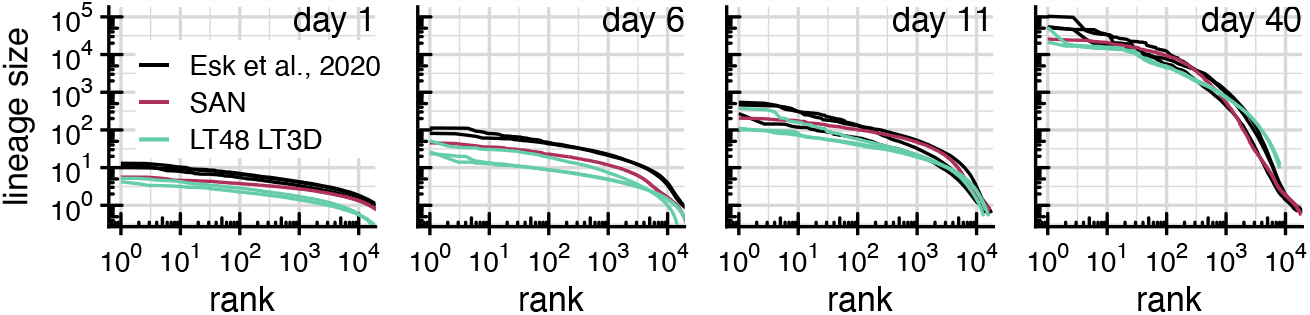
Replicate experiments. Replicate experiments based on the same organoid protocol show similar lineage size distributions as the data from Esk et al. (2020). Ranks of the Esk et al. data and SAN model predictions were scaled to account for an 1.7-fold increase in the number of detected lineages in the replicate experiments.

**Figure S5.**
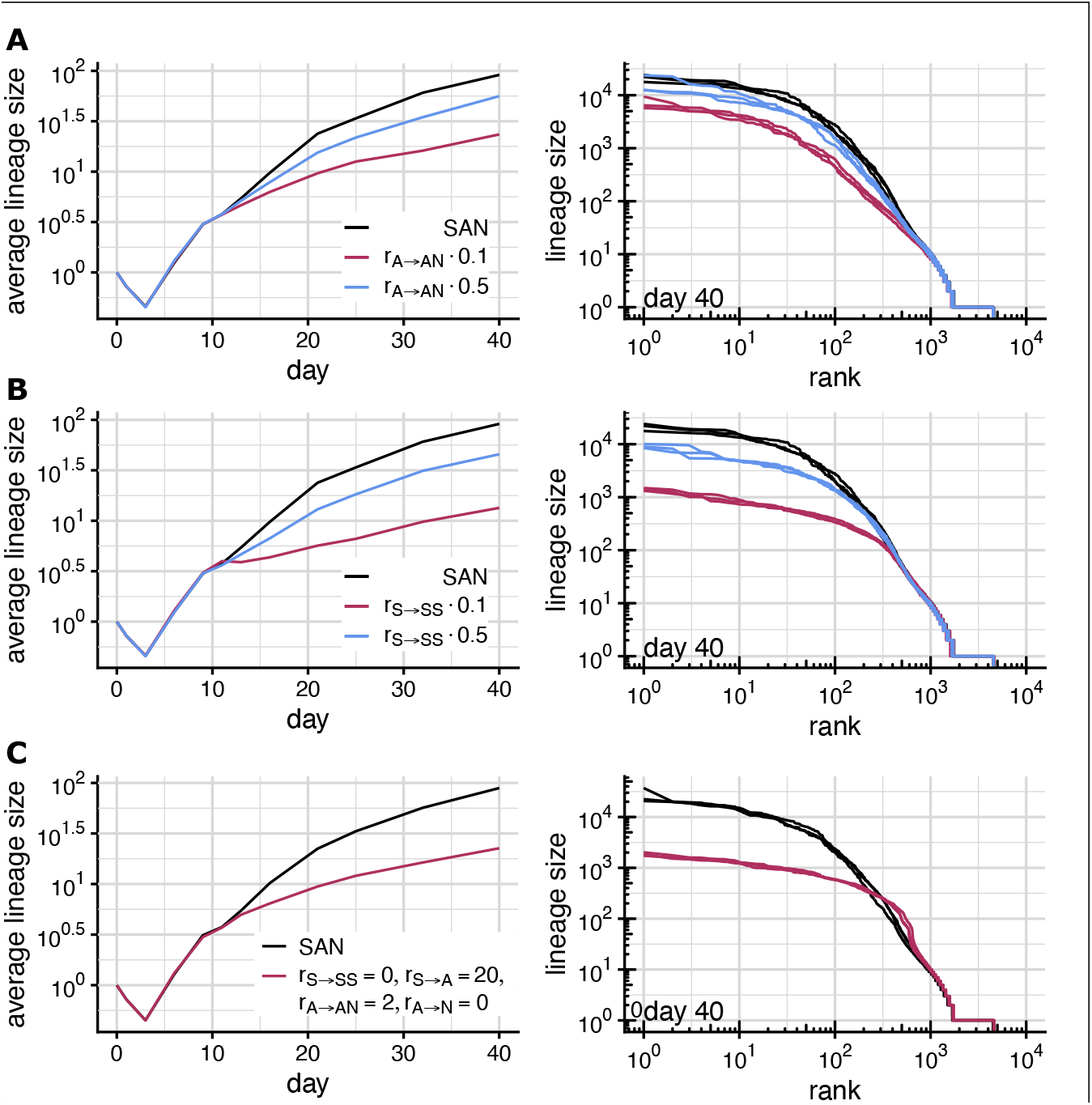
Perturbation Simulations. **(A)**. Rate of asymmetric division after day 11 reduced to one-half (blue) and one-tenth (black) of its original value. **(B)**. Rate of symmetric division after day 11 reduced to one-half (blue) and one-tenth (black) of its original value. Rate of S -> A differentiations reduced accordingly to keep the net S-cell growth rate unchanged **(C)**. Immediate differentiation of S-cells on day 11, no symmetric divisions.

